# Single dose of amphetamine induces delayed subregional attenuation of striatal cholinergic interneuron activity

**DOI:** 10.1101/2021.03.22.436417

**Authors:** Samira Ztaou, Soo Jung Oh, Sophia Tepler, Sixtine Fleury, Miriam Matamales, Jesus Bertran-Gonzalez, Nao Chuhma, Stephen Rayport

**Affiliations:** Department of Molecular Therapeutics, New York State Psychiatric Institute, New York, NY 10032, USA; Department of Psychiatry, Columbia University, New York, NY 10032, USA; Decision Neuroscience Laboratory, School of Psychology, University of New South Wales, Sydney, NSW, Australia

**Keywords:** Psychostimulant, Dopamine, Acetylcholine, Fluorescence Imaging, Phosphorylated ribosomal protein S6

## Abstract

Psychostimulants such as amphetamine target dopamine neuron synapses to engender drug-induced plasticity. While dopamine neurons modulate the activity of striatal cholinergic interneurons (ChIs) with regional heterogeneity, how amphetamine affects ChI activity has not been elucidated. Here, we applied quantitative fluorescence imaging approaches to map the dose-dependent effects of a single dose of amphetamine on ChI activity at 2.5 and 24 hours after injection across the mouse striatum using the activity-dependent marker phosphorylated ribosomal protein S6 (p-rpS6). We found that amphetamine did not induce neurotoxic effects on ChIs, as their distribution and morphology were not affected. While low- or high-dose amphetamine did not affect ChI activity after 2.5 hours, ChI activity was significantly reduced in all striatal subregions after 24 hours, with a dose-dependent effect in the nucleus accumbens. Thus, our findings suggest that a single dose of amphetamine has delayed regionally heterogeneous effects on ChI activity.

**Significance statement:** Using the activity dependent marker phosphorylated ribosomal protein S6 (p-rpS6), we mapped amphetamine effects on the activity of cholinergic interneurons (ChIs) across the striatum. Amphetamine caused a delayed attenuation of ChI activity in all striatal subregions, and a dose-dependent effect in the ventral striatum/nucleus accumbens, a critical site of psychostimulant action.

## Introduction

Psychostimulants such as amphetamine (AMPH) target DA neuron terminals (Pierce and Kalivas, 1997; Luscher and Malenka, 2011; Sulzer, 2011) and engender dose-dependent behavioral effects. DA release in the ventral Str/nucleus accumbens (NAc) is associated with hyperlocomotion whereas DA release in the dorsal Str is associated with stereotypies (Robinson and Becker, 1986; Kalivas and Stewart, 1991; Gaytan et al., 1998; Yates et al., 2007). DA neurons modulate the activity of cholinergic interneurons (ChIs), which comprise less than 2% of striatal (Str) neurons, and yet strongly control the Str circuitry (Goldberg and Wilson, 2010; Gonzales and Smith, 2015; Abudukeyoumu et al., 2019). Modulation of ChI activity is critical in processing and reinforcement of reward-related behaviors (Atallah et al., 2014; Gonzales and Smith, 2015). ChIs in the ventral Str are crucial for psychostimulant-dependent behaviors (Sofuoglu and Mooney, 2009; Witten et al., 2010; Lee et al., 2020; Lewis and Borrelli, 2020). However, whether the effects of AMPH on ChIs occur at specific striatal loci has not been elucidated.

Several studies have previously shown that the phosphorylated form of the ribosomal protein S6 (p-rpS6), a ubiquitous translational activation marker, can be used to estimate the cellular activity of ChIs under different pharmacological and/or behavioral conditions (Bertran-Gonzalez et al., 2012; Matamales et al., 2016a, 2016b). Pharmacological silencing or increasing ChI firing lead to a striking decrease or increase of p-rpS6 signal in ChIs, respectively (Bertran-Gonzalez et al., 2012; Matamales et al., 2016b). Unlike several other activity-dependent markers (Chandra and Lobo, 2017), p-rpS6 is predominantly expressed in ChIs (Bertran-Gonzalez et al., 2012). The phosphorylation of rpS6 can be induced by multiple signaling cascades; mTORC1 pathway and/or mTORC1-independent pathways such as the PKC, the MAPK or the cAMP/PKA pathways (Valjent et al., 2011; Bertran-Gonzalez et al., 2012; Gangarossa and Valjent, 2012). The phosphorylation of rpS6 appears to occur sequentially at 5 serine residues: in the order 236, 235, 240, 244 and 247 (Knight et al., 2012; Biever et al., 2015a). In particular, the phosphorylation of rpS6 at serine 240 and 244 residues (p-rpS6^240/244^) is a reporter of ChI activity.

To address regionality in AMPH modulation of ChI activity in the striatum, we mapped p-rpS6 intensity in ChIs throughout the entire rostrocaudal axis after a single low- or high-dose of AMPH at two time points. This revealed that AMPH induces a delayed regionally heterogeneous dose-dependent attenuation of Str ChI activity.

## Materials and Methods

### Ethics

This research was performed in accordance with the Guide for the Care and Use of Laboratory Animals of the National Institutes of Health, under a protocol approved by the Institutional Animal Care and Use Committee of New York State Psychiatric Institute (#NYSPI-1494).

### Experimental animals

Sixteen male and fourteen female DAT-IRES-Cre/+;ROSA26-flox-STOP-CAG-ChR2-YFP double mutant mice (Jackson Laboratories, RRID:IMSR_JAX:006660, RRID:IMSR_JAX:024109), the same genotype as previous studies (Chuhma et al., 2014, 2018; Mingote et al., 2015, 2017), were used at postnatal day (P) 56-82. Mice were 129Sv/C57BL6J mixed background, backcrossed to C57BL6J at least 5 times and kept inbred. Mice were maintained on a 12:12 hours dark/light cycle with lights on at 7:00 AM in a temperature-controlled room with food and water provided *ad libitum*.

### Drug treatment

D-amphetamine hemisulfate (Sigma-Aldrich, A5880) either low-dose (2 mg/kg) or high-dose (16 mg/kg) was dissolved in 0.9% NaCl immediately before use. Injections were done intraperitoneally (i.p.) at a volume of 10 ml/kg body weight.

### Behavioral monitoring

Mice were habituated to handling for 2 days prior to the drug administration. Monitoring took place under bright ambient light conditions during the light phase. On the injection day, mice were placed in the open field, equipped with infrared motion detectors (Plexiglas activity chambers, 40.6 cm long × 40.6 cm wide × 38.1 cm high; SmartFrame Open Field System, Kinder Scientific) for 1 hour for habituation. Baseline activity was monitored for 30 min pre-injection, then mice were injected i.p. either with saline, 2 mg/kg or 16 mg/kg AMPH, and observed for a 2 hour post-injection period. Locomotor activity was recorded automatically in 10-min bins. Stereotyped behaviors − orofacial stereotypy (mouth movements, lick, bite, self-gnaw, taffy pull, jaw tremor, yawn) and grooming − were scored for 1 min every 5 min as previously described (Kelley, 2001). One low-dose AMPH-injected mouse, in the 2.5 hours post-injection (2.5h_pi_) cohort, was excluded from the study as its locomotor activity decreased after injection.

### Immunocytochemistry

For immunocytochemistry, mice were deeply anesthetized with ketamine (90 mg/kg)/xylazine (7 mg/kg) and then perfused intracardially with cold PBS (100 mM; pH 7.4) followed by 4% paraformaldehyde (PFA). Brains were removed and post-fixed overnight in 4% PFA. Coronal sections were cut at 50 μm with a vibrating microtome (Leica VT1200S), and stored in a cryoprotectant solution (30% glycerol, 30% ethylene glycol in 0.1 M Tris HCl, pH 7.4) at −20 °C. Free-floating sections were washed in PBS and incubated in glycine (100 mM) for 30 min to quench aldehydes. Non-specific binding was blocked with 10% normal donkey serum (NDS) in 0.1 PBS Triton X-100 for 2 hours. Sections were incubated in PBS containing 0.02% Triton X-100 and 2% NDS overnight at 4°C with primary antibodies: anti-ChAT (1:500, goat polyclonal, Millipore Cat# AB144P, RRID:AB_2079751) and anti-phosphorylated ribosomal protein S6 (p-rpS6^240/244^, 1:1500, rabbit polyclonal, Cell Signaling Technology Cat# 2215, RRID:AB_331682). Sections were then washed with PBS, and secondary antibodies applied for 45 min in PBS containing 0.02% Triton X-100 at room temperature: anti-goat Alexa Fluor 594 (1:200, Thermo Fisher Scientific Cat# A-11058, RRID:AB_2534105) and anti-rabbit Alexa Fluor 488 (1:200, Thermo Fisher Scientific Cat# A-21206, RRID:AB_2535792). Sections were mounted on gelatin subbed slides (SouthernBiotech) and cover slipped with ProLong Gold aqueous medium with DAPI (Thermo Fisher Scientific) and stored at 4°C until imaging.

### Imaging and Analysis

Images were acquired using an Axio Imager M2 fluorescence microscope (Zeiss) with a high-resolution digital camera (Axiocam 506 mono, 2752 × 2208 pixels, Zeiss), a 20×/0.8 objective and Zen 2.3 Digital Imaging software (Zeiss; RRID:SCR_013672). Ten coronal sections, spanning the rostrocaudal extent of the right Str (bregma 1.54, 1.18, 0.98, 0.62, 0.26, −0.10, −0.46, −0.82, −1.22 and −1.58 mm), were imaged. An image stack consisting of 5 planes at 5 μm intervals was obtained. Exposure time for each excitation was held constant throughout acquisition.

Raw 16-bit images were analysed using Fiji/ImageJ (Version 2.0.0., NIH, RRID:SCR_002285). Z-projected images were obtained by taking pixels with the maximum intensity in a stack. The outer boundary of the Str and its anatomical subregions − nucleus accumbens core (NAc core) and shell (NAc shell), dorsomedial (DM Str) and dorsolateral (DL Str) Str − were manually delineated in accordance with the mouse brain atlas (Paxinos and Franklin, 2008), and their areas (mm^2^) in each coronal section were obtained.

Particle analysis detected all ChAT-positive neurons in the Str and the total number of ChIs, perimeter (μm), area (μm^2^), and circularity (a circularity value of 1 indicates a perfect circle while values approaching 0 indicate more elongated shapes) of each ChI were measured. Density of ChIs (neurons/mm^2^) in each Str subregion was calculated as ChI number in a subregion divided by area of the subregion. For each coronal section, the ChAT image was superimposed on the p-rpS6 image, and the ChAT-positive neurons were used as a mask for p-rpS6 intensity analysis. Fluorescence intensity of the corpus callosum was used for background subtraction.

Location of each ChI was defined by coordinates of the centroid. The p-rpS6 intensity of each ChI was normalized to the maximum and the minimum intensities for each cohort − 2.5h_pi_ and 24h_pi_ − and color-scaled. All color-scaled ChIs were 3D plotted with outlines of the Str using a customized script in MATLAB (MathWorks; RRID:SCR_001622) as previously described (Matamales et al., 2016a).

Distributions of p-rpS6 fluorescence intensity were standardized to the corresponding saline group for each time point and subregion by calculating z-scores: z = (x-μ)/σ, where x is the p-rpS6 signal in individual ChI, μ and σ are the mean and the standard deviation, respectively, of p-rpS6 signal in the corresponding saline group.

### Statistical analysis

Sample size estimation was done with G*Power 3.1 (G*Power, RRID: SCR_013726), setting α = 0.05 and power = 0.8 (Cunningham and McCrum-Gardner, 2007). Effect size was estimated from previous experiments (Cohen’s D = 0.97), giving a required n per group of 5. Statistical analyses were performed using Prism 8 (GraphPad Prism, RRID:SCR_002798) or SPSS 26 (SPSS; RRID:SCR_002865). P < 0.05 was considered as significant for all analyses. Data are presented as mean ± SEM.

Parametric tests were used here because datasets followed a normal distribution (D’Agostino-Pearson normality test, p > 0.05). ANOVA was used for comparison among conditions. Where significance was detected, multiple pairwise comparison with Bonferroni correction was performed as a *post-hoc* test.

## Results

### Dose-dependent effects of AMPH on locomotor activity and stereotyped behavior

Mice were studied after a single low- (2 mg/kg) or high-dose (16 mg/kg) of AMPH at 2.5 hours post-injection (2.5h_pi_), after acute behavioral effects have subsided, and at 24h_pi_ to assess enduring effects on Str activity. To confirm differential behavioral effects of the two AMPH doses, mice received saline, low-dose or high-dose AMPH, and their locomotion and stereotypy were monitored for 2 hours in the open field (**Fig. 1A**). Total travel distance dose-dependently increased in both the 2.5h_pi_ cohort (saline 17.6 ± 3.4 m, low-dose 94.5 ± 13.8 m, high-dose 212.5 ± 29.3 m) and 24h_pi_ cohort (saline 18.6 ± 3.4 m, low-dose 64.0 ± 5.0 m, high-dose 302.7 ± 35.8 m), however no significant difference was observed between the two time points (two-way ANOVA; treatment effect, *F*_(2, 24)_ = 78.15, *p* < 0.001; time effect, *F*_(1, 24)_ = 1.55, *p* = 0.23) (**Fig. 1B**, **left**). Although there was a significant treatment × time interaction (*F*_(2, 24)_ = 4.95, *p* = 0.02), these two cohorts showed similar dose-dependent hyperlocomotion, a significant increase after low-dose and a further increase after high-dose.

**Figure 1.**
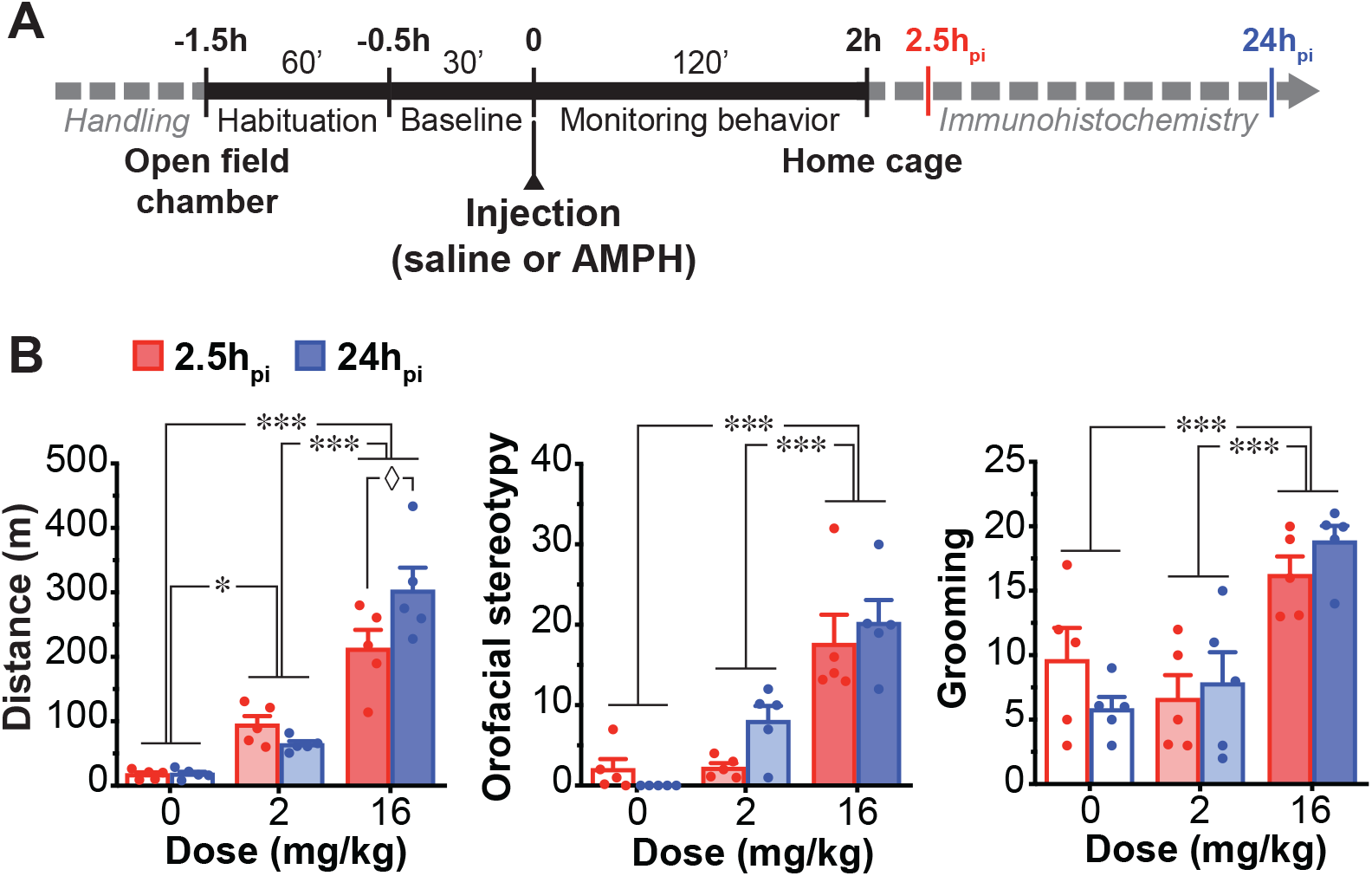
Effects of a single dose of AMPH on behavior. (**A**) Timeline of amphetamine (AMPH) experiments. (**B**) Total traveled distance (left), orofacial stereotypy (middle) and grooming (right) score after saline (0 mg/kg, n = 5 animals), low-dose (2 mg/kg, n = 5 animals) or high-dose (16 mg/kg, n = 5 animals) AMPH, at 2.5 hours post-injection (2.5h_pi_) (red) and 24h_pi_ (blue). Dots in bar graphs show measurements from each animal. *, ** and *** indicate p < 0.05, p < 0.01 and p < 0.001, respectively; ◊ indicates p < 0.05 for comparison between 2.5h_pi_ and 24h_pi_.

Low- and high-dose AMPH increased orofacial stereotypy in both 2.5h_pi_ cohort (saline 2.0 ± 1.3, low-dose 2.2 ± 0.6, high-dose 17.6 ± 3.6) and 24h_pi_ cohort (saline 0 ± 0, low-dose 8.0 ± 1.9, high-dose 20.2 ± 2.9), however no significant difference was observed between the two time points (two-way ANOVA; treatment effect, *F*_(2, 24)_ = 38.74, *p* < 0.001; time effect, *F*_(1, 24)_ = 1.50, *p* = 0.23; treatment × time interaction, *F*_(2, 24)_ = 1.69, *p* = 0.21) (**Fig. 1B**, **middle**).

Low-dose AMPH did not affect grooming score in the two cohorts while high-dose increased it in both 2.5h_pi_ cohort (saline 9.6 ± 2.5, low-dose 6.6 ± 1.9, high-dose 16.2 ± 1.5) and 24h_pi_ cohort (saline 5.8 ± 0.9, low-dose 7.8 ± 2.4, high-dose 18.8 ± 1.2) (two-way ANOVA; treatment effect, *F*_(2, 24)_ = 19.86, *p* < 0.001). Neither a significant difference nor a treatment × time interaction was observed between the two time points (time effect, *F*_(1, 24)_ = 0, *p* > 0.99; treatment × time interaction, *F*_(2, 24)_ = 1.67, *p* = 0.21) (**Fig. 1B**, **right**). These observations confirmed that AMPH showed a comparable dose-dependent behavioral activation in the two cohorts, 2.5h_pi_ and 24h_pi_.

### Distribution and morphology of ChIs is not altered by AMPH

To address potential neurotoxic effects of AMPH on ChIs (Zhu et al., 2006 607), we examined the distribution of ChIs as well as their soma morphology. ChIs were identified by ChAT immunostaining and examined in 10 coronal sections spanning the rostrocaudal extent of the right Str in four subregions: nucleus accumbens (NAc) core and shell, dorsomedial (DM) and dorsolateral (DL) Str (**Fig. 2A**). The previously recognized rostro-caudal distribution of ChIs (Matamales et al., 2016a) peaked at 0.98 mm from bregma and gradually declined caudally. The distribution was not affected by either AMPH dose or time after injection (**Fig. 2B**) (three-way ANOVA; rostrocaudal effect, *F*_(9, 240)_ =204.13, *p* < 0.001; treatment effect, *F*_(2, 240)_ = 0.17, *p* = 0.85; time effect, *F*_(1, 240)_ = 0.66, *p* = 0.42; rostrocaudal × treatment × time interaction, *F*_(18, 240)_ = 1.43, *p* = 0.12). Although numbers of ChIs varied significantly between Str subregions, AMPH or time after injection did not affect ChI count significantly in any Str subregion (three-way ANOVA; location effect, *F*_(3, 96)_ = 954.82, *p* < 0.001; treatment effect, *F*_(2, 96)_ = 0.12, *p* = 0.89; time effect, *F*_(1, 96)_ = 0.42, *p* = 0.52; location × treatment × time interaction, *F*_(6, 96)_ = 0.52, *p* = 0.80) (**Fig. 2C**). Although ChI densities varied between Str subregions, highest in the NAc shell and lowest in the NAc core, AMPH did not affect densities in any subregion or time point (three-way ANOVA; location effect, *F*_(3, 96)_ = 73.76, *p* < 0.001; treatment effect, *F*_(2, 96)_ = 0.15, *p* = 0.86; time effect, *F*_(1, 96)_ = 0.57, *p* = 0.45; location × treatment × time interaction, *F*_(6, 96)_ = 0.30, *p* = 0.94) (**Fig. 2D**).

**Figure 2.**
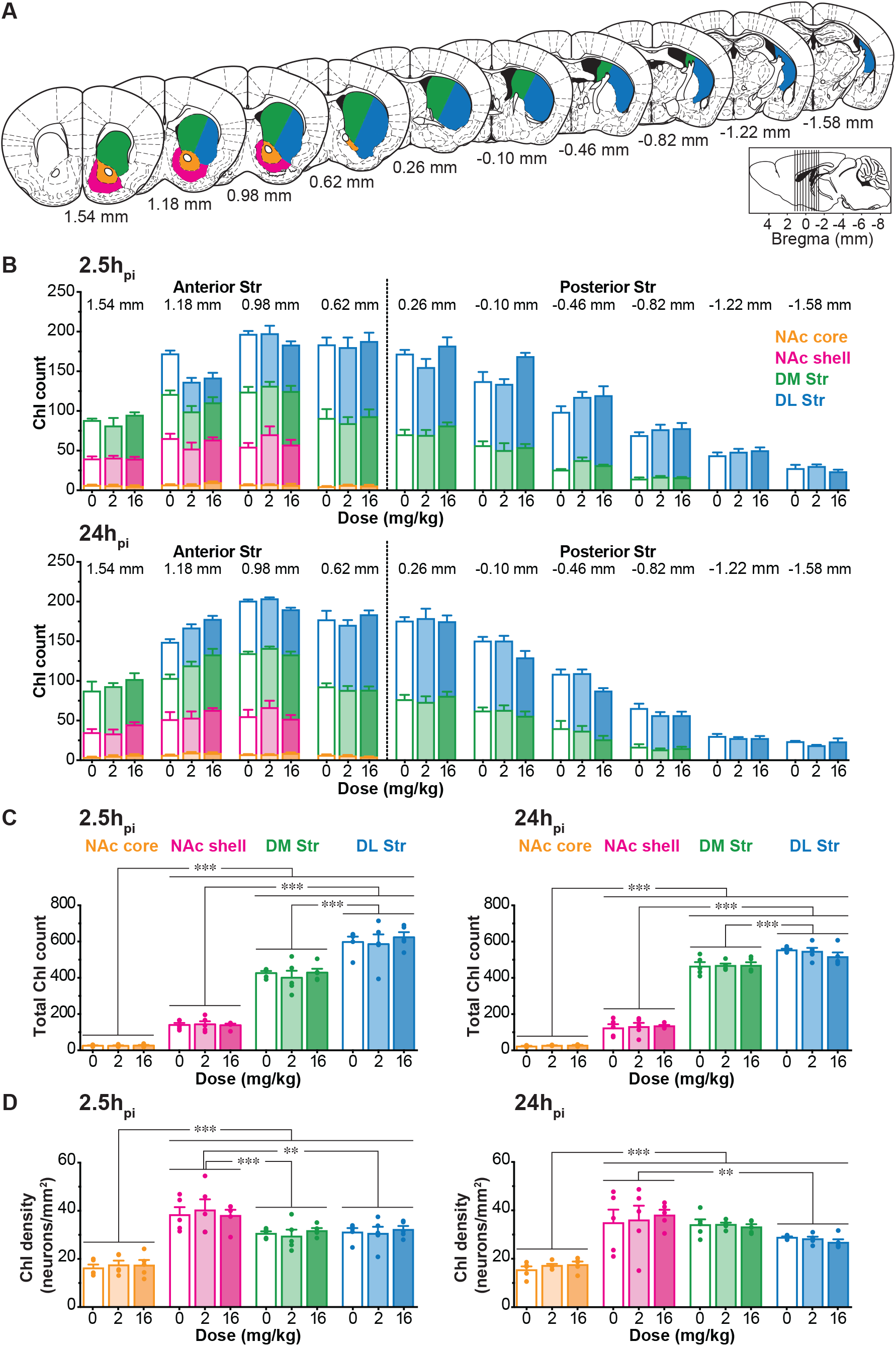
Distribution of ChIs 2.5h_pi_ and 24h_pi_ AMPH. (**A**) Schematic of 10 coronal sections of the striatum (Str) (from bregma +1.54 mm to −1.58 mm). Delineations of Str subregions are shown in the right Str: nucleus accumbens core (NAc core, orange), nucleus accumbens shell (NAc shell, magenta), dorsomedial Str (DM Str, green), dorsolateral Str (DL Str, blue). Locations of slices are shown in inset, and locations from bregma are indicated under the slices. (**B**) Stacked bar graphs showing counts of ChIs across Str subregions (NAc core, orange bar; NAc shell, magenta bar; DM Str, green bar; DL Str, blue bar) in 10 coronal hemisections along the rostrocaudal axis after saline (0 mg/kg, n = 5 animals), low-dose (2 mg/kg, n = 5 animals) or high-dose (16 mg/kg, n = 5 animals) AMPH, at 2.5h_pi_ (top) and 24h_pi_ (bottom). (**C**) Total ChI count in each Str subregion, at 2.5h_pi_ (left) and 24h_pi_ (right). (**D**) ChI density (neurons/mm^2^) in each Str subregion, at 2.5h_pi_ (left) and 24h_pi_ (right). Dots in bar graphs show the average measurements from each animal. ** and *** indicate p < 0.01 and p < 0.001, respectively.

AMPH did not affect ChI soma area (2.5h_pi_: saline 242.5 ± 4.4 μm^2^, low-dose 243.6 ± 10.5 μm^2^, high-dose 240.4 ± 9.6 μm^2^; 24h_pi_: saline 242.6 ± 14.6 μm^2^, low-dose 250.1 ± 3.0 μm^2^, high-dose 242.5 ± 5.5 μm^2^; two-way ANOVA; treatment effect, *F*_(2, 24)_ = 0.21, *p* = 0.81; time effect, *F*_(1, 24)_ = 0.16, *p* = 0.69; treatment × time interaction, *F*_(2, 24)_ = 0.07, *p* = 0.94), perimeter (2.5h_pi_: saline 72.3 ± 1.4 μm, low-dose 74.3 ± 3.4 μm, high-dose 75.3 ± 3.1 μm; 24h_pi_: saline 76.3 ± 1.1 μm, low-dose 75.6 ± 1.4 μm, high-dose 76.3 ± 1.4 μm; two-way ANOVA; treatment effect, *F*_(2, 24)_ = 0.23, *p* = 0.80; time effect, *F*_(1, 24)_ = 1.44, *p* = 0.24; treatment × time interaction, *F*_(2, 24)_ = 0.29, *p* = 0.75) or circularity (2.5h_pi_: saline 0.60 ± 0.02, low-dose 0.59 ± 0.04, high-dose 0.56 ± 0.03; 24h_pi_: saline 0.56 ± 0.04, low-dose 0.58 ± 0.04, high-dose 0.55 ± 0.04; two-way ANOVA; treatment effect, *F*_(2, 24)_ = 0.28, *p* = 0.76; time effect, *F*_(1, 24)_ = 0.56, *p* = 0.46; treatment × time interaction, *F*_(2, 24)_ = 0.07, *p* = 0.93) in the whole Str, at any time points (**Fig. 3A, B**).

**Figure 3.**
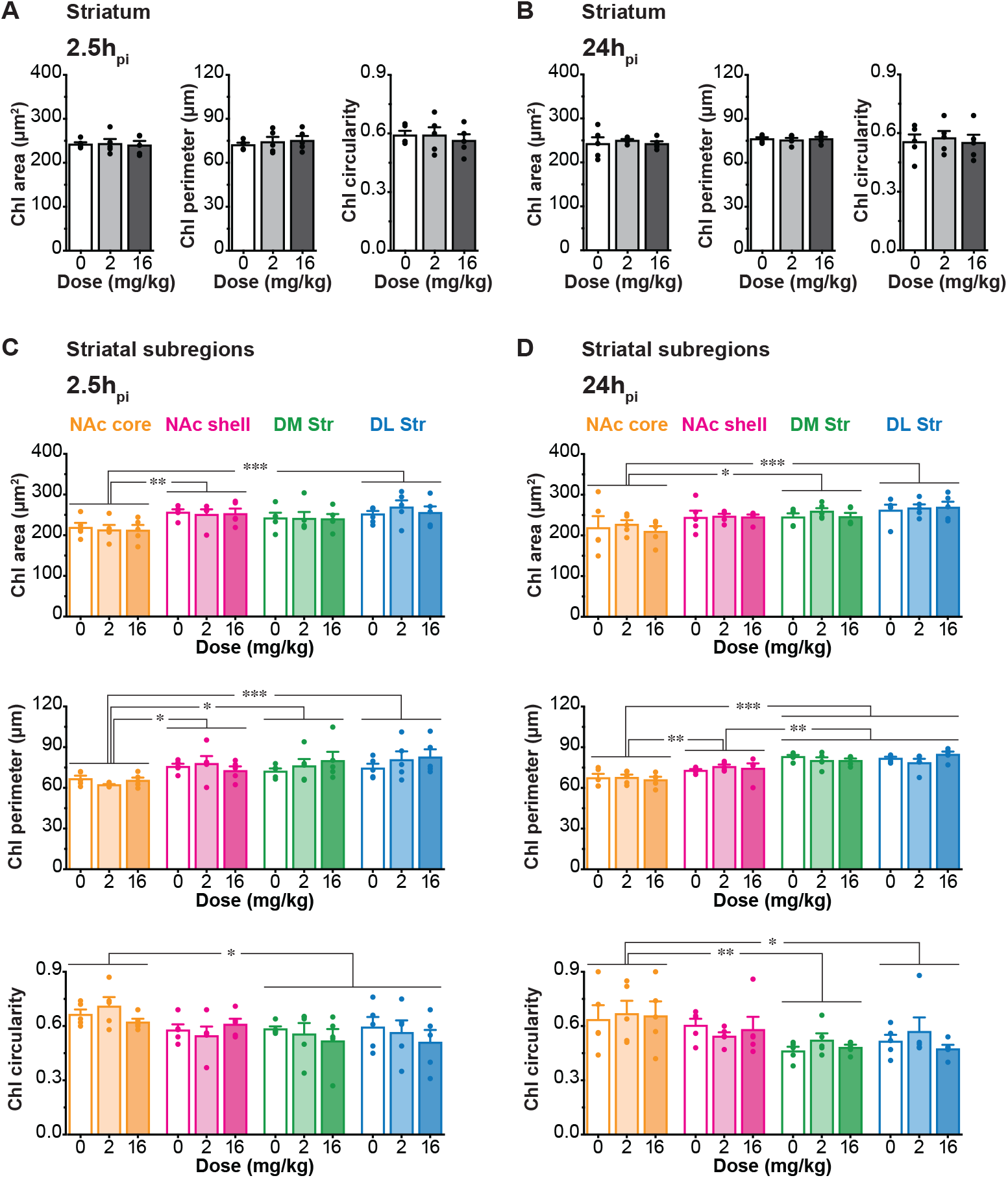
Morphology of ChIs 2.5h_pi_ and 24h_pi_ AMPH. (**A**-**B**) Morphological characteristics of ChIs: area (μm^2^), perimeter (μm) and circularity in 10 coronal hemisections in the whole Str after saline (0 mg/kg, white bar, n = 5 animals), low-dose (2 mg/kg, gray bar, n = 5 animals) or high-dose (16 mg/kg, black bar, n = 5 animals) AMPH, at 2.5h_pi_ (**A**) and 24h_pi_ (**B**). (**C**-**D**) Area (top), perimeter (middle) and circularity (bottom) of ChIs in each Str subregion (NAc core, orange bar; NAc shell, magenta bar; DM Str, green bar; DL Str, blue bar) at 2.5h_pi_ (**C**) and 24h_pi_ (**D**). Dots in bar graphs show the average measurements from each animal. *, ** and *** indicate p < 0.05, p < 0.01 and p < 0.001, respectively.

Although ChI morphology differed between Str subregions, AMPH did not affect soma area, perimeter, or circularity in any Str subregion or at the two different time points (area: three-way ANOVA; location effect, *F*_(3, 96)_ = 13.17, *p* < 0.001; treatment effect, *F*_(2, 96)_ = 0.37, *p* = 0.68; time effect, *F*_(1, 96)_ = 0.31, *p* = 0.58; location × treatment × time interaction, *F*_(6, 96)_ = 0.18, *p* = 0.98; perimeter: three-way ANOVA; location effect, *F*_(3, 96)_ = 22.45, *p* < 0.001; treatment effect, *F*_(2, 96)_ = 0.38, *p* = 0.68; time effect, *F*_(1, 96)_ = 2.42, *p* = 0.12; location × treatment × time, *F*_(6, 96)_ = 0.83, *p* = 0.55; circularity: three-way ANOVA; location effect, *F*_(3, 96)_ = 7.69, *p* < 0.001; treatment effect, *F*_(2, 96)_ = 0.58, *p* = 0.57; time effect, *F*_(1, 96)_ = 1.36, *p* = 0.25; location × treatment × time interaction, *F*_(6, 96)_ = 0.35, *p* = 0.91) (**Fig. 3C, D**). Thus, neither low- nor high-dose AMPH affected ChI morphology, arguing against neurotoxic effects of a single dose of AMPH.

### AMPH attenuated ChI activity across Str at 24h_pi_

We mapped AMPH effects on ChI activity with p-rpS6 staining. Double immunostaining showed colocalization of ChAT and p-rpS6 (**Fig. 4**), although p-rpS6 signal was also detected in other Str neurons (Bertran-Gonzalez et al., 2012). We quantified p-rpS6 intensity as the average pixel intensity in each ChAT positive neuron, in saline, low- and high-dose AMPH-injected mice, at 2.5h_pi_ or 24h_pi_ (n = 5 animals/condition, 10 hemisections/animal). Individual ChI locations were plotted in coronal hemisections of the Str and p-rpS6 intensities were color-scaled (**Fig. 5A, C**).

**Figure 4.**
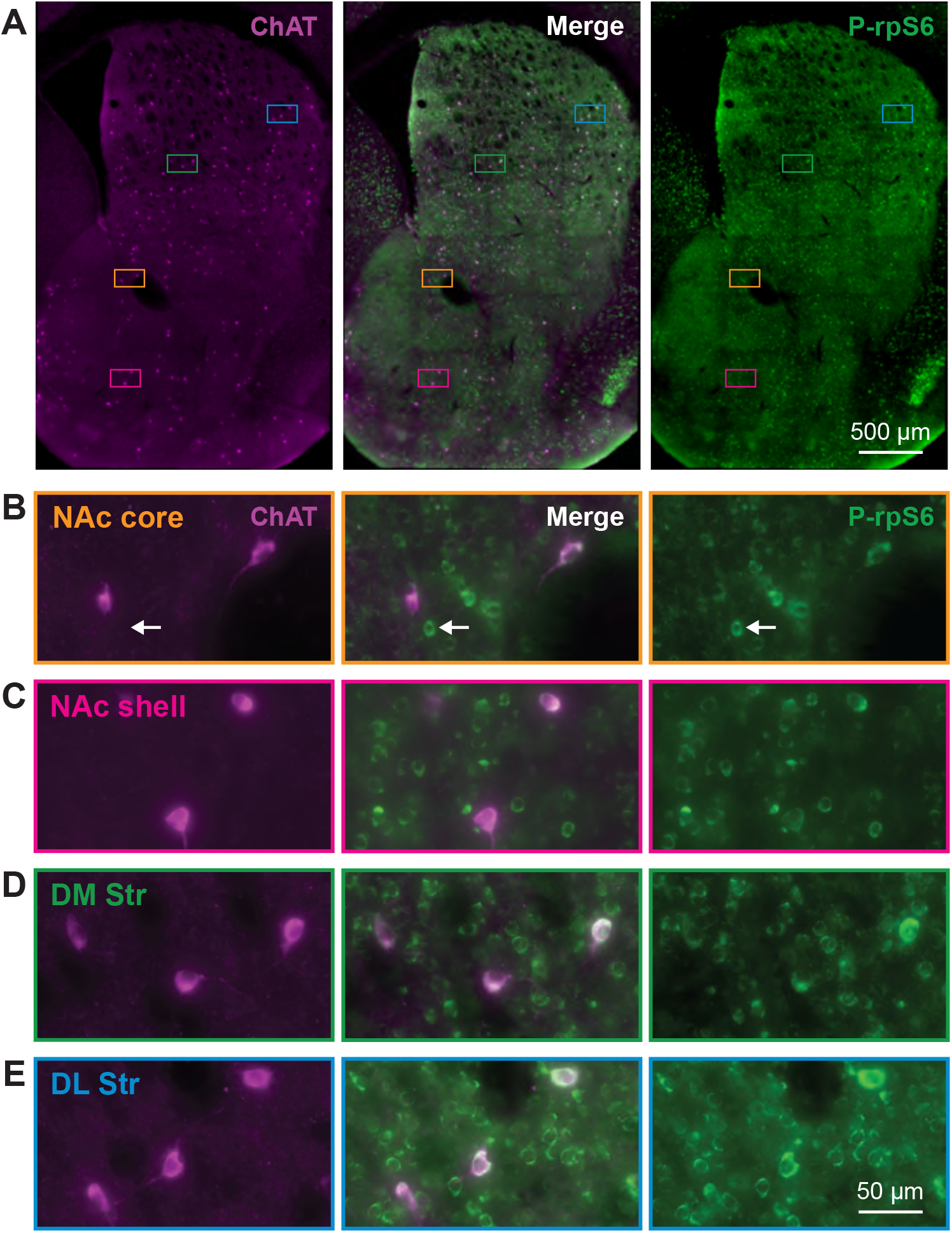
Phosphorylation of ribosomal protein S6 in ChIs. (**A**) Low magnification images of ChAT (purple) and p-rpS6 (green) in a Str section (bregma 0.98 mm) with merged images in the middle. Colored rectangles are representative locations of Str subregions and magnified in B-D. Expanded images of the NAc core (**B**, orange), NAc shell (**C**, magenta), DM Str (**D**, green) and DL Str (**E**, blue) subregions. Arrows indicate a ChAT-negative p-rpS6-positive neuron.

**Figure 5.**
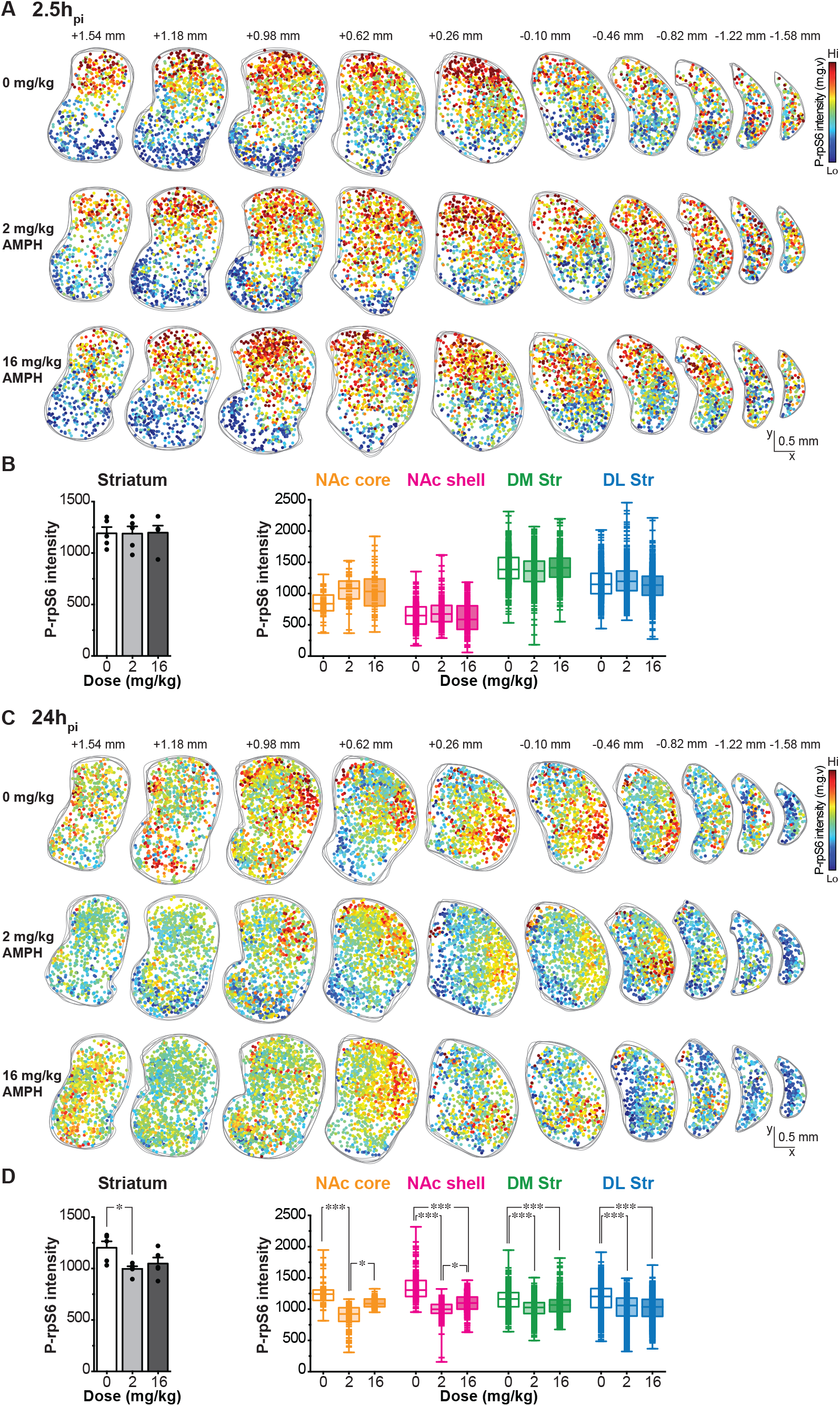
P-rpS6 intensity in ChIs 2.5h_pi_ and 24h_pi_ AMPH. (**A**-**C**) Spatial distribution of ChIs with relative p-rpS6 intensity shown in 10 coronal hemisections along the rostrocaudal axis (from bregma +1.54 mm to −1.58 mm) at 2.5h_pi_ (**A**) and 24h_pi_ (**C**). The spatial distribution of ChIs from five animals was superimposed for each injection group: saline (top), low-dose (middle) or high-dose (bottom) AMPH. Each dot represents one ChI and intensity of p-rpS6 is shown by a blue (low level) to red (high level) color scale. (**B**-**D**) Left: averaged p-rpS6 intensity in ChIs in the whole Str at 2.5h_pi_ (**B**) or 24h_pi_ (**D**) of either saline (0 mg/kg, white bar, n = 5 animals), low-dose (2 mg/kg, gray bar, n = 5 animals) or high-dose (16 mg/kg, black bar, n = 5 animals) AMPH. Each dot in bar graphs indicates the averaged p-rpS6 intensity from each animal. Right: Box and whiskers plots showing p-rpS6 intensity in ChIs in each Str subregion (NAc core, orange bar; NAc shell, magenta bar; DM Str, green bar; DL Str, blue bar) at 2.5h_pi_ (**B**) and 24h_pi_ (**D**). Each line symbol in bar graphs shows the p-rpS6 intensity in individual ChI. * and *** indicate p < 0.05 and p < 0.001, respectively.

At 2.5h_pi_, p-rpS6 intensity varied among Str subregions, with higher p-rpS6 intensity in the DM Str decreasing ventrally to the NAc shell (**Fig. 5A**). There was no apparent difference in the distribution of p-rpS6 intensity between saline, low-dose, and high-dose AMPH-injected animals (**Fig. 5A**). Average p-rpS6 intensity in the whole Str did not differ between saline, low- and high-dose AMPH-injected animals (one-way ANOVA, *F*_(2, 12)_ = 0.003, *p* = 0.99) (**Fig. 5B**, **left**). Although average ChI p-rpS6 intensities differed between Str subregions, neither low- nor high-dose AMPH affected ChI p-rpS6 intensity in any Str subregion at 2.5h_pi_ (two-way ANOVA; location effect, *F*_(3, 3503)_ = 246.30, *p* < 0.001; treatment effect, *F*_(2, 3503)_ = 1.85, *p* = 0.16; location × treatment interaction, *F*_(6, 3503)_ = 1.89, *p* = 0.08) (**Fig. 5B**, **right**).

At 24h_pi_, ChI p-rpS6 intensity was reduced by low-dose AMPH in all Str regions, particularly the NAc (**Fig. 5C**). Low-dose AMPH reduced p-rpS6 staining more than high-dose (**Fig. 5C**). Indeed, low-dose AMPH reduced average ChI p-rpS6 staining in the whole Str, while high-dose AMPH did not show a significant effect (one-way ANOVA, *F*_(2, 12)_ = 4.35, *p* = 0.03) (**Fig. 5D**, **left**). Low-dose AMPH reduced ChI p-rpS6 intensity in all Str subregions, while high-dose reduced the intensity in the NAc shell, DM Str, and DL Str, but not in the NAc core (**Fig. 5D**, **right**) (two-way ANOVA; location effect, *F*_(3, 4543)_ = 4.19, *p* = 0.006; treatment effect, *F*(2, 4543) = 62.05, *p* < 0.001; location × treatment interaction, *F*_(6, 4543)_ = 3.51, *p* = 0.002). In the NAc core and shell, low-dose AMPH reduced ChI activity more than high-dose, while the two doses caused a similar reduction in the dorsal Str.

To compare AMPH effects on p-rps6 intensity between the two time points, p-rpS6 intensities in ChIs were compared to the respective saline groups by z-scores for each Str subregion. Z-scores in AMPH-injected animals at 2.5h_pi_ did not differ significantly from saline-injected in any Str subregion (**Fig. 6A**) (two-way ANOVA; treatment effect *F*_(2, 3503)_ = 2.12, *p* = 0.12; location effect *F*_(3, 3503)_ = 4.49, *p* = 0.004; treatment × location interaction *F*_(6, 3503)_ = 2.01, *p* = 0.06). At 24h_pi_, p-rpS6 intensity z-scores became negative after low- or high-dose AMPH in all Str subregions, indicating a reduction in ChI activity (**Fig. 6B**) (two-way ANOVA; treatment effect, *F*_(2, 3503)_ = 76.78, *p* < 0.001; location effect, *F*(3, 3503) = 4.39, *p* = 0.004; treatment × location interaction *F*_(6, 4543)_ = 4.65, *p* < 0.001). AMPH effects on z-scores at 24h_pi_ were significantly different from those at 2.5h_pi_ (three-way ANOVA; time effect, *F*_(1, 96)_ = 27.35, *p* < 0.001; treatment effect, *F*_(2, 96)_ = 3.18, *p* = 0.04; location effect, *F*_(3, 96)_ = 0.34, *p* = 0.80; time × treatment × location interaction, *F*_(6, 96)_ = 0.89, *p* = 0.51).

**Figure 6.**
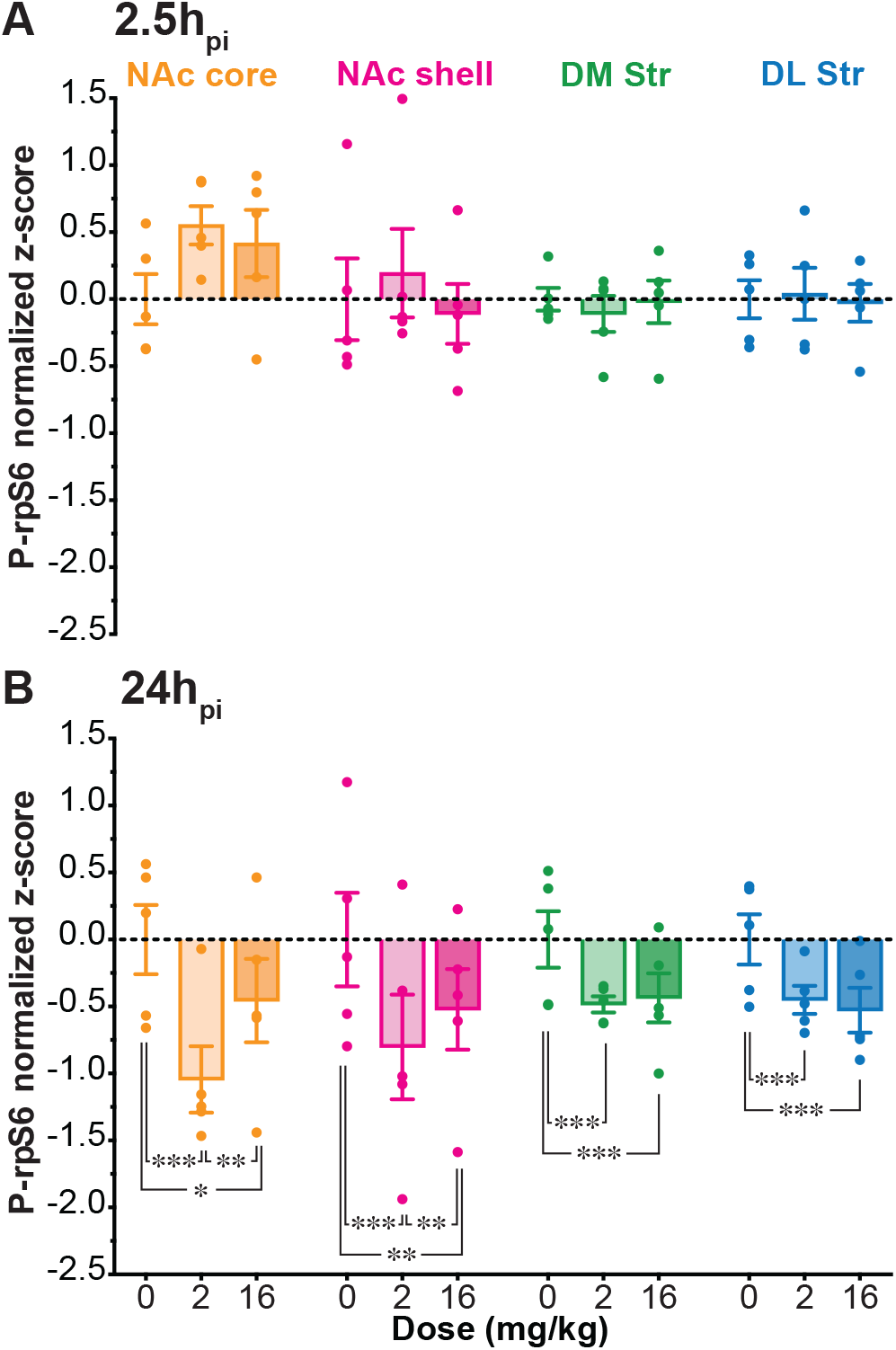
Comparison of p-rpS6 intensity in ChIs. (**A**-**B**) Comparison of ChI p-rpS6 intensity z-score normalized to mean intensity in corresponding saline-injected animals 2.5h_pi_ (**A**) and 24h_pi_ (**B**) of saline (0 mg/kg, n = 5 animals), low-dose (2 mg/kg, n = 5 animals) or high-dose (16 mg/kg, n = 5 animals) AMPH in each Str subregion (NAc core, orange bar; NAc shell, magenta bar; DM Str, green bar; DL Str, blue bar). Dots in bar graphs show the average measurements from each animal. *, **, and *** indicate p < 0.05, p < 0.01, and p < 0.001, respectively.

As in the raw intensity data at 24h_pi_ (**Fig. 5**), in the ventral subregions, the Nac core showed a similar profile of attenuation to the Nac shell. Likewise, in the dorsal subregions, the DM Str showed a similar profile of attenuation to the DL Str. This suggests a high degree of replication in the ventral and dorsal Str, respectively. At 24h_pi_, in the NAc core and shell, low-dose had more of an effect than high-dose, while the effects of the two doses were not significantly different in the dorsal Str (**Fig. 6B**). Thus, the difference between the 2.5h_pi_ and 24h_pi_ cohorts reflect a time- and subregion dependent effect of a single dose of AMPH on ChI activity in the Str.

## Discussion

Str ChIs are principal targets of the striatal DA innervation and subject to regionally heterogeneous modulation. Here, we mapped the downstream effects of a single AMPH dose on ChI activity using p-rpS6 as a ChI-selective activity-dependent marker. Importantly, the single dose of AMPH did not affect the distribution or overall morphology of ChIs in any Str subregion. While AMPH had no effect on ChI activity at 2.5h_pi_, AMPH significantly attenuated ChI activity at 24h_pi_ in all subregions. In the NAc, the reduction in ChI activity after low-dose was less than after high-dose. In the dorsal Str, low- and high-dose AMPH produced a smaller attenuation. Thus, a single dose of AMPH significantly reduces ChI activity throughout the Str at 24h_pi_, with a dose-dependency in the NAc but not the dorsal Str.

### Distribution, morphology and activity of ChIs in the Str

We confirmed the differential distribution of ChIs in striatal subregions (Gonzales and Smith, 2015) and that this distribution as well as the morphology of ChIs were not affected by either low- or high-dose AMPH, at 2.5h_pi_ or 24h_pi_. ChIs are denser in the NAc medial shell, as previously described in mice (Matamales et al., 2016a), rats (Phelps and Vaughn, 1986), and primates (Brauer et al., 2000). The ChAT-positive neuropil is also denser (Phelps and Vaughn, 1986; Brauer et al., 2000).

In rodents (Gonzales and Smith, 2015), non-human primates (Brauer et al., 2000) and humans (Holt et al., 1996), average sizes of ChIs in the NAc are smaller than those in the dorsal Str. Here, we found that ChIs in the NAc core were significantly smaller and more elongated compared to those in other Str subregions, and that the morphology of ChIs soma (area, perimeter and circularity) differed between Str subregions. A single injection of AMPH, either low- or high-dose, did not affect ChI distribution or soma morphology in any Str subregion, at either time point, showing these doses did not induce neurotoxic effects on ChIs. Although AMPH neurotoxicity on DA has been known for some time (Wagner et al., 1980; Ricaurte et al., 1984; Ryan et al., 1990; Miller and O’Callaghan, 1996; Krasnova et al., 2001; Krasnova et al., 2005; Granado et al., 2018), no study focused on AMPH downstream neurotoxic effect on Str ChIs. To cause a significant toxic effect on ChIs, a higher dose of a more potent psychostimulant appears to be required; a single high-dose (30 mg/kg) of methamphetamine was found to induce a loss of 29% of ChIs in the dorsal Str (Zhu et al., 2006).

The present results confirm the dorsoventral gradient in ChI activity (Matamales et al., 2016a). P-rpS6 signal reports the integrated activity and its modulation appears to be detected 60 min after pharmacological or behavioral manipulations (Bertran-Gonzalez et al., 2012), suggesting that p-rpS6 is suitable to study ChI activity at 2.5h_pi_ or later (Knight et al., 2012). Therefore, the lack of AMPH effect at 2.5h_pi_ is not due to detection limitations of p-rpS6 measurement. Stress increases P-rpS6 intensity (Knight et al., 2012; Biever et al., 2015a) as seen in the greater intensity 2.5h_pi_ compared to 24h_pi_ in the saline controls.

### Single dose of AMPH affects ChI activity

P-rpS6 intensity in ChIs was not affected at 2.5h_pi_ after low- or high-dose AMPH, while ChI activity modulation via DA neuron glutamatergic cotransmission is dose-dependently attenuated after a single dose of AMPH at 2.5h_pi_ (Chuhma et al., 2014). This discrepancy could be due to differences in the measurements; p-rpS6 reflects the tonic *in vivo* activity of ChIs, which receive cortical and thalamic inputs in addition to DA neuron inputs, compared to the short phasic firing control of ChIs by DA neuron synaptic inputs.

In a previous study, we demonstrated that a single dose of AMPH at 2.5h_pi_ dose-dependently attenuates the inhibitory DA D2 receptor action on ChIs (Chuhma et al., 2014). Clinical studies have shown that chronic psychostimulant use like cocaine, methamphetamine and AMPH is associated with an overall downregulation of DA transmission, both DA release and D2 receptor levels (Ashok et al., 2017; Proebstl et al., 2019). Here, we could have expected an increase in ChI activity due to the loss of D2 receptor inhibition. ChIs are also targets of glutamate cotransmission of DA neurons in addition to subsecond D2 receptor mediated firing inhibition (Chuhma et al., 2014, 2018; Mingote et al., 2019). The lack of AMPH effect on ChI activity at 2.5h_pi_ could be due to reduction in both glutamate cotransmission and DA.

The attenuation of ChI activity at 24h_pi_ is likely to be polysynaptic effects rather than direct effects on DA neuron presynaptic terminals. ChIs receive cortical and particularly strong thalamic glutamatergic inputs (Lim et al., 2014). Tonic attenuation of cortical or thalamic glutamatergic inputs may be caused by polysynaptic modulation, resulting in delayed attenuation of ChI activity. 2.5 hours appear not to be sufficient to cause long-term circuit change, since AMPH does not affect p-rpS6^240/244^ level or protein synthesis in the Str within 2 hours following injection (Rapanelli et al., 2014; Biever et al., 2015b).

In the present study, low-dose AMPH reduced ChI activity most in the ventral Str/NAc, a crucial site of psychostimulant action (Russo et al., 2010; Sulzer, 2011), while high-dose affected all Str subregions to the same extent. Behavioral effects of high-dose AMPH, induced both hyperlocomotion and stereotypies. These regional effects are in line with hyperlocomotion mediated by ventral tegmental area DA system projecting to NAc and stereotypies are mediated by the substantia nigra DA system projecting to the dorsal Str (Kuczenski, 1983; Kuczenski et al., 1991).

### ChIs in psychostimulant-dependent behavior

DA neurons projecting to the ventral Str/NAc corelease glutamate (Hnasko et al., 2010; Stuber et al., 2010) and can drive burst firing in ChIs (Chuhma et al., 2014; Mingote et al., 2019). A single dose of AMPH attenuates glutamate cotransmission (Chuhma et al., 2014), and mice with conditional reduction in glutamate cotransmission show an attenuated sensitization to repeated AMPH (Mingote et al., 2017), suggesting that DA neuron connections to ChIs in the NAc medial shell regulate AMPH responsiveness. Here, we have found that low-dose AMPH particularly affects ChIs in the ventral Str/NAc further supporting the critical role for DA neuron glutamate cotransmission to ChIs.

Although psychostimulant abuse is characterized by repeated use, a single dose of AMPH can induce enduring Str circuit changes, drug-dependent behavior and negative affective states, such as anhedonia, depression and anxiety (Vanderschuren et al., 1999; Koob and Le Moal, 2001; Xia et al., 2008; Kameda et al., 2011; Li et al., 2017; Jing et al., 2018; Rincon-Cortes et al., 2018; Jayanthi et al., 2020). Our results, in line with these previous findings, point to the relevance of a single dose of AMPH to elucidate drug-induced plasticity. Enduring alterations in ChI activity following acute AMPH exposure point to this neuronal population as a key component in mediating AMPH effects in the Str circuitry.

## Acknowledgments

We thank Susana Mingote, Leora Yetnikoff and Vlad Velicu for technical help and advice. This work was supported by NIH R01 DA038966 and R01 MH117128 (SR), Philippe Foundation (SZ) and by ARC DP190102511, ARC DP210102700 and NHMRC APP1165990 (JBG and MM) and FT200100502 (JBG).

